# Celastrol sensitizes bicalutamide to enhance immunogenic Cell death in the treatment of castration-resistant prostate

**DOI:** 10.1101/2025.07.08.663785

**Authors:** Peipei Meng, He Li, Yuhang Meng, Dong Li, Guancheng Wang, Qinxin Zhao, Weibin Yang, Huang Hongdong

## Abstract

**Background:** Prostate cancer is the most commonly diagnosed non-cutaneous malignancy in men and is the second-leading cause of cancer-related death. Castration-resistant prostate cancer (CRPC) persists despite low testosterone levels.

**Methods:** In this study, we investigated the combined effect of Celastrol and bicalutamide on prostate cancer cells. We evaluated cytotoxicity, apoptosis, oxidative stress, ferro ptosis and the activation of the immune system *in vivo* and *in vitro*.

**Results:** Our findings revealed that the co-administration of Celastrol with bicalutamide enhanced cytotoxicity against prostate cancer cells, promoted cell apoptosis, induced oxi dative stress and ferroptosis. Importantly, it assisted bicalutamide in activating the immune system, triggering immunogenic cell death (ICD), and thereby enhancing immune therapy. This synergy resulted in a stronger anti-CRPC effect, improving the overall therapeutic efficacy.

**Conclusion:** Our study demonstrates the potential of the Celastrol and bicalutamide combination in treating castration-resistant prostate cancer. Further studies are warranted to explore the clinical application of this approach and its potential in improving patient outcomes.

**Funding information:** The author(s) declare that financial support was received for the research and/or publication of this article. This work was supported by Beijing Municipal Natural Science Foundation (grant no. BFHHQS20240003).

## 1. Introduction

Prostate cancer (Pca) is one of the deadliest cancers affecting men globally. It is the third most common malignant tumor and the fourth leading cause of cancer-related deaths in men (1-3), and is particularly prevalent in older men over 65 years of age. In 202 2, there will be an estimated 1.47 million new cases of prostate cancer worldwide, and 397,000 deaths due to prostate cancer (3). Prostate cancer often presents with subtle symptoms, leading to many patients being diagnosed at advanced stages. This results in a lack of early detection and frequent bone metastases at diagnosis (4-6). After systematic diagnosis and assessment of prostate cancer, treatment options include radical surgery, radical radiotherapy, chemotherapy, endocrine therapy, and others. Among these, androgen deprivation therapy (ADT) is the standard of care for the initial management of advanced or metastatic prostate cancer (7), but progression to castration-resistant prostate cancer (CR PC) occurs within 2-3 years of initiation of ADT. Currently, there are limited therapeutic means to treat CRPC, and the existing therapeutic regimens are not satisfactory. Consequently, there is an immediate necessity to investigate new therapeutic strategies to enhance survival outcomes for patients with CRPC.

Studies concluded that CRPC remains an androgen-dependent disease and that deprivation levels should be maintained throughout its course (8-11). bicalutamide, the first-gener ation antiandrogen, is used in PCA medication (12). It is combined with luteinizing horm one-releasing hormone (LHRH) agents in patients with metastatic hormone-sensitive prostate cancer to block testosterone flare. Due to its high target selectivity, bicalutamide generally does not interact with other steroid hormone receptors and therefore does not have o ther off-target hormonal activity. bicalutamide is active at the androgen receptor and competes directly with androgens (e.g., testosterone, dihydrotestosterone), preventing their binding and inhibiting androgen agonism, thus completely removing androgenic activity. Although bicalutamide lacks androgenic inhibitory capacity, it also does not exert antigonadotro pic or androgen synthesis inhibitory effects. Instead, it achieves its anti-androgenic effect by blocking androgens at the receptor. Prostate cancer cells are prone to developing drug resistance after long-term treatment with bicalutamide. This resistance results in mutations in the androgen receptor and increased expression of AR splice variants, which hinder effective drug binding to the receptor. Therefore, the development of drugs that improve the sensitivity of bicalutamide to the receptor is necessary.

Celastrol (Cel) is an active alkaloid extracted from either the whole plant or the peeled woody parts of the traditional Chinese medicine Lei Gong Teng (Tripterygium wilfordii Hook. f). Celastrol can activate both endogenous and exogenous apoptotic pathways, leading cancer cells to undergo apoptosis. This includes increasing the expression of the pr o-apoptotic protein Bax and decreasing the expression of the anti-apoptotic protein Bcl-2. Recent studies have shown that Celastrol significantly inhibits the growth of various types of tumor cells (13-15). Celastrol in combination with Olaparib was found to reduce prostate cancer survival and migration (16). Additionally, Celastrol has been found to prom ote DNA damage and apoptosis in other tumor types while also enhancing immune responses (17-19). We found that bicalutamide together with Celastrol, strengthens the body’s anti-tumor immune response.

Ferroptosis is an iron-dependent form of Cell death (20). It is characterized by iron accumulation and uncontrolled lipid peroxidation, which leads to plasma membrane rupture and the release of intracellular contents (21). Lipid peroxidation is a central factor in triggering ferroptosis. Growing evidence indicates that ferroptosis significantly contributes t o tumorigenesis, progression, metastasis, and drug resistance. Resistance to ferroptosis is a common feature of tumor cells. Elevated levels of ROS associated with this resistance can induce mutations in tumor genes and enhance the malignancy of these cells, ultimately leading to drug resistance (22, 23). Tumor cells can develop drug resistance through t he SLC7A11-GSH-GPX4 axis. Down-regulating SLC7A11 can induce ferroptosis, decrease GSH synthesis, and inactivate GPX4, which inhibits tumor cell growth (24).

This paper will explore the mechanisms of action of Celastrol and bicalutamide, focusing on their effects on prostate cancer cell proliferation, iron death induction, invasion a nd metastasis inhibition, and in vivo tumor immunotherapy modulation. Ultimately, the a alysis of Celastrol — a natural compound with therapeutic efficacy — aims to garner m ore attention from the scientific and pharmacological communities and expedite its clinical application for the benefit of cancer patients. This article aims to provide a new perspective on CRPCs and discuss their role in facilitating the search for new drugs.

## 2. Methods

All chemicals were obtained from commercial sources and used without further purification unless otherwise noted. Cel [(2R,4aS,6aS,12bR,14aS,14bR)-10-hydroxy-2,4a,6a,9,12b, 14a-hexamethyl-11-oxo-1,2,3,4,4a,5,6,6a,11,12b,13,14,14a,14b-tetradecahydropicene-2-carboxylic acid] was purchased from Bide Pharm (Shanghai, China). RPMI 1640 media, Dulbecco’s Modified Eagle Medium/Nutrient Mixture F-12 (DMEM/F12), fetal bovine serum (FB S), and trypsin were purchased from Gibco (BRL, MD, USA). Whole protein extraction kit, 4′,6-diamidino-2-phenylindole (DAPI), and MTT were purchased from Solarbio (Beijing, China). ROS assay kit and ATP assay kit were purchase from Beyotime (Shanghai, China). Live & dead viability/cytotoxicity assay kit were purchase from KeyGEN biotechn ology (Nanjing, China). Annexin V–FITC/PI apoptosis detection kit, Calcein/PI Cell viability/cytotoxicity assay kit, and 4% paraformaldehyde were purchased from Beyotime (Shanghai, China).

### 2.1 General instruments

Flow cytometry (FCM) was conducted by Cytomics FC500 Flow Cytometry (Beckman Coulter, U.S.A.). Confocal laser scanning microscopy (CLSM) was accomplished by ZEISS LSM880. All OD values were recorded by Bio-Tek.

### 2.2 Cell lines and animals

The PC-3, 22RV1, C4-2, and RM-1 Cells were preserved in the Beijing National Laboratory for Molecular Sciences, State Key Laboratory of Polymer Physics and Chemistry, Institute of Chemistry (Beijing, China). C57BL/6 mice were purchased from SPF Biotechnology (Beijing, China). The mice were maintained in an environment with a temperature of 22 ± 1°C, a relative humidity of 50 ± 1%, a light/dark cycle of 12 hours/12 hour s, and provided with sterile food and water. All animal experiments were carried out under the guidelines evaluated and approved by the ethics committee of Beijing Friendship Hospital, Capital Medical University, China (MR-11-24-028093).

### 2.3 IntraCellular ROS Generation

The levels of intraCellular ROS were evaluated using the H2DCF-DA (2’,7’-dichlorodihydrofluorescein diacetate) assay. For the experiment, PC-3 Cells were first treated with PBS, bicalutamide (Bica), Celastrol (Cel), or a combination of Bica and Cel for 24 hours. After treatment, all Cells were incubated with 10μM H2DCF-DA for 30 minutes, allowing for ROS detection. Following incubation, the Cells were washed twice with PBS to remove excess dye. For nuclear visualization, Cells were then stained with DAPI for 10 minutes. Fluorescence images were captured using CLSM, and the intraCellular ROS production was quantified using FCM.

### 2.4 Cell viability assays

The PC-3, 22RV1, C4-2, and RM-1 cell lines were seeded in 96-well plates at a density of 8 × 10^3^ cells per well and allowed to adhere for 12 hours under standard culture conditions. Following initial attachment, cells were exposed to experimental treatments consisting of DMSO (vehicle control), Bicalutamide (Bica), or Celastrol (Cel) at concentrations ranging from 0.125 to 100 μM. After 48 hours of drug exposure, cellular viability was assessed using the MTT assay. Briefly, 10% MTT reagent (3-(4,5-dimethylthiazol-2-yl)-2,5-diphenyltetrazolium bromide) was added to each well and incubated for 4 hours at 37°C. The resulting formazan crystals were subsequently solubilized by adding 100 μl o f SDS solubilization solution (10% sodium dodecyl sulfate in 0.01 M HCl) to each well, followed by an overnight incubation. Optical density measurements were obtained using a BioTek microplate reader with dual-wavelength detection at 570 nm (primary absorbance) and 650 nm (reference wavelength).

### 2.5 Apoptosis analysis

Apoptosis analysis was performed using the Annexin V-FITC/PI apoptosis detection k it (Beyotime, C1052) following standardized protocols. In brief, PC-3 cells were plated in 12-well culture plates at a density of 2 × 10^5^ cells/well and allowed to adhere overnight. Following 12-hour incubation under standard culture conditions (37°C, 5% CO2), cells were subjected to experimental treatments comprising: (1) PBS (vehicle control), (2) Bicalutamide (Bica), (3) Celastrol (Cel), and (4) Bica + Cel combination therapy for 24 hour s. Post-treatment, cells were gently washed twice with ice-cold PBS and subsequently sta ined with Annexin V-FITC/propidium iodide (PI) working solution (prepared per manufacturer’s specifications) for 30 minutes at 37°C under light-protected conditions. Cellular flu orescence was immediately quantified using flow cytometry with excitation at 488 nm an d emission detection at 530 nm (FITC) and 575 nm (PI). Data analysis was performed using FlowJo software, with apoptotic populations quantified as early apoptotic (Annexin V?/PI?) and late apoptotic/necrotic (Annexin V?/PI?) cells.

### 2.6 Measurement of the Release of HMGB1

PC-3 Cells were seeded in glass-bottomed culture dishes at a density of 1.0 × 10^5^ C ells per well and incubated for 12 hours. After incubation, the Cells were treated with P BS, Bica, Cel, or Bica + Cel for 24 hours. Following the treatment, the Cells were was hed with PBS and fixed in 4% paraformaldehyde for 15 minutes at room temperature. T he Cells were then permeabilized with 0.1% Triton X-100 for 10 minutes and blocked with 1% BSA for 30 minutes. After blocking, the Cells were incubated overnight at 4°C with the HMGB1 primary antibody diluted in blocking buffer. The next day, the Cells w ere washed with PBS and incubated with an Alexa Fluor 555-conjugated secondary antib ody for 1 hour at room temperature. Finally, the nuclei were stained with DAPI for 10 minutes, and the samples were imaged using a CLSM to examine HMGB1 expression an d localization.

### 2.7 Measurement of Cell Surface CRT

PC-3 Cells (1.0×10^5^) were seeded onto cover slides in a 24-well plate and incubated at 37°C for 12 hours. Cells were then treated with PBS, Bica, Cel, Bica + Cel for 24 hours. After treatment, Cells were washed with PBS, fixed in 4% paraformaldehyde solution, and permeabilized with 0.1% Triton X-100. Cells were blocked with 1% BSA for 2 0 minutes, then incubated overnight with a diluted CRT primary antibody. The following day, Cells were washed and incubated with an Alexa Fluor 555-conjugated secondary antibody for 2 hours. Nuclear DNA was stained with DAPI, and the samples were imaged using a CLSM to assess CRT exposure on the Cell surface.

### 2.8 ATP release assay

ExtraCellular ATP levels were measured using an ATP assay kit (Beyotime, S0026), following the manufacturer’s protocol. PC-3 Cells were seeded in 12-well plates at a dens ity of 2 × 10^5^ Cells per well and allowed to adhere overnight. After incubation, the Cell s were treated with PBS, Bica, Cel, and Bica + Cel for 24 hours. The culture medium was then collected and centrifuged at 1000 rpm for 3 minutes to remove Cell debris. 10 0 μL of ATP detection reagent was added to each well and incubated at room temperature for 5 minutes. Subsequently, 20 μL of the supernatant or standard solutions were added, and the luminescence was measured using a microplate reader to quantify ATP concentration.

### 2.9 BMDC mature evaluation *in vitro*

Bone marrow-derived dendritic Cells (BMDCs) were isolated from 4–6-week-old male C57BL/6 mice. The Cells were cultured in DMEM medium supplemented with 10% FB S, 20 ng/mL granulocyte-macrophage colony-stimulating factor (GM-CSF), and 10 ng/mL interleukin-4 (IL-4), and maintained at 37°C with 5% CO2. After a 3-5 day culture period, the BMDCs were co-cultured with pre-treated RM-1 Cells at a 1:1 ratio for an additional 24 hours. Following incubation, the BMDCs were harvested, washed with PBS, and stained with the following antibodies: anti-CD11c-PE, anti-CD80-FITC, and anti-CD86-A PC, for 1 hour at room temperature. The Cells were then analyzed by FCM to assess dendritic Cell maturation markers.

### 2.10 *In vivo* antitumor efficacy evaluation

RM-1 prostate cancer cells (8 × 10^5^ cells in 100 μL Matrigel/PBS mixture) were orthotopically implanted into the right posterior flank of 6-week male C57BL/6 mice (n=5/group) under aseptic conditions. Following tumor engraftment, mice were randomly stratified into four treatment cohorts using block randomization: (1) PBS (vehicle control, daily oral gavage), (2) Bicalutamide (Bica, 50 mg/kg, oral gavage every other day), (3) Celastrol (Cel, 0.15 mg/kg, intraperitoneal injection every other day), and (4) Bica + Cel com bination therapy. Treatments commenced on day 0 post-implantation and continued through day 12 with alternating administration schedules to prevent drug interaction.

Tumor progression was monitored triweekly using digital caliper measurements, with volume calculated as (length × width^2^)/2. Body weight and clinical signs were recorded daily to assess treatment toxicity. On day 14, mice were humanely euthanized via CO2 asphyxiation followed by cervical dislocation. Excised tumors were weighed using analytical balances, measured in three dimensions, and documented with standardized photographic protocols.

### 2.11 Immunohistochemical and immunofluorescence analyses

The solid tumor were harvested from RM-1 tumor-bearing mice on the 14th day of tumor inoculation for histological observation by standard H&E staining, immunohistochem ical and immunofluorescence staining. For H&E staining, the excised tumor and organs were fixed in 4% paraformaldehyde solution, embedded in paraffin, sectioned, and stained with H&E. The sections were then observed under a fluorescence microscope. For detecting the expression of CRT, HMGB1 and infiltration of CD8^+^ T Cells in tumor tissues, frozen tumor sections were fixed, and blocked with 1% BSA. Then the sections were incubated with primary antibodies against CRT, HMGB1 and CD8 overnight at 4 °C, followed by processing with corresponding second antibodies. Nuclei were counterstained with DAPI and then the stained sections were imaged with a CLSM.

### 2.12 Flow cytometric analysis of tumor immune microenvironment

Fresh tumors, spleen, and draining lymph node tissue were collected for antitumor immune response analysis via FACS. Samples were briefly dissociated into single-Cell sus pensions, followed by removal of red blood Cells by red blood Cell lysis buffer (Solabio). Subsequently, the samples were blocked with 0.1% BSA in PBS and incubated with relevant antibodies at room temperature for 1 h. To characterize T Cells, TAM in the tumor, the Cells underwent staining with anti-mouse CD3-PE, anti-mouse CD4-APC, anti-mouse CD8-FITC, anti-mouse F4/80-PE, anti-mouse CD80-FITC, and anti-mouse CD206-AP

C. To analyze T Cells in spleen, Cells were stained with anti-mouse CD3-PE, anti-mouse CD4-APC, anti-mouse CD8-FITC. For the analysis of DCs in tumors and lymph nodes, Cells were stained with anti-mouse CD11c-PE, anti-mouse CD80-FITC, and anti-mouse CD86-APC. Flow cytometric data acquisition was performed with CytExpert software, an d the data were processed with FlowJo software.

### 2.13 Western blotting

PC-3 cells were incubated with various pharmacological agents for 24 hours at 37°C. Then the cells were lysed using RIPA lysis buffer supplemented with 1 mM phenylmethane-sulfonyl fluoride (PMSF). The cells were lysed using ultrasound for 30 minutes at 4° C, then centrifuged at 12,000 rpm for 15 minutes to collect the supernatant. The protein concentration was quantified using a BCA protein concentration assay kit (Beyotime, P0 011), after which SDS-PAGE buffer was added. Subsequently, the protein lysates were denatured in a metal bath at 95°C and subjected to SDS-polyacrylamide gel electrophoresis (SDS-PAGE) for separation. The separated proteins were then transferred onto a PVDF membrane (Merck Millipore) and blocked using skim milk for 2 hours at room temperature. After blocking, the membrane was incubated overnight at 4°C with the following primary antibodies at a dilution of 1:1000: GPX4 (Beyotime, AF7020), SCD (Beyotime, AF 7944), SLC7A11 (Beyotime, AF7992) and β-Tubulin (Beyotime, AF1216). The PVDF membrane was subsequently washed (5 min × 5) with tris-buffered saline with tween (TBS T) buffer and then incubated with the appropriate secondary antibody (Peroxidase-conjugated goat anti-rabbit IgG (H + L), at a dilution of 1:1000) in a 5% bovine serum albumin solution for 2 h at 25°C. After further washing with TBST buffer, membrane was photographed using the chemiluminescent imaging system.

### 2.14 Statistical analysis

Statistical analyses were conducted using GraphPad Prism 9 software. Data are expressed as mean ± SD from a minimum of three independent biological replicates, unless otherwise indicated in the figure legends. The significance between groups was determined by unpaired two-sided t-tests, one-way ANOVA with Dunnett’s multiple comparisons, or t wo-way ANOVA with Tukey’s multiple comparisons, as appropriate. A p-value of 0.05 or less was considered statistically significant. The significance levels were denoted as follows: p < 0.05, **p < 0.01, ***p < 0.001, and **** p < 0.0001.

## 3. Results

### 3.1 In vitro antitumor activity study

First, the anticancer activity of Celastrol was investigated in four prostate cancer cell lines: PC-3, 22RV1, C4-2, and RM-1. MTT assay results showed that the half-maximal inhibitory concentration (IC50) of Celastrol was highest in RM-1 Cells, at 14.01μM, while it was lowest in PC-3 Cells, at 10.13μM (Figure 1A). Additionally, the IC50 values in 22RV1 and C4-2 Cells were 6.047μM and 17.91μM, respectively (Figure 1A). The MT T results indicated no significant difference in tumor cell death among the four prostate cancer cell lines treated with bicalutamide. In this study, the PC-3 cell line was selected as the model for in vitro experiments, using a bicalutamide dose of 25μM.

Next, we analyzed apoptosis in PC-3 cells treated with different agents using Annexin V-FITC and propidium iodide (PI) double staining. The results showed that the apoptosis rate was highest in cells treated with the combination of Celastrol and bicalutamide a t 40.54%, which was 2.06 times higher than in cells treated with bicalutamide (9.39%) (Figures 1B, 1C). Then, live-dead staining was performed to observe the distribution of live and dead Cells in PC-3 Cells. In cells treated with Celastrol and bicalutamide, we observed strong red fluorescence. This indicates a significantly higher number of dead cell s compared to those treated with only bicalutamide (Figure 1D). Finally, DNA in the Cel ls was stained using Edu to observe how active the Cell proliferation was. It was observed by CLSM that the green fluorescence in the Cells treated with Celastrol and bicalutamide, which would have been fluorescently labeled with EdU, was weaker than that in the control and bicalutamide-treated groups, suggesting that the proliferative activity of the tumor Cells was reduced by the Celastrol and bicalutamide treatment (Figure 1F). FCM analysis revealed that EdU fluorescence values in cells treated with bicalutamide were 1. 07 times higher than those treated with Celastrol and 1.09 times higher than in cells trea ted with both Celastrol and bicalutamide (Figure 1E). In conclusion, Celastrol significantly enhances the antitumor activity of bicalutamide *in vitro*.

**Fig. 1.**
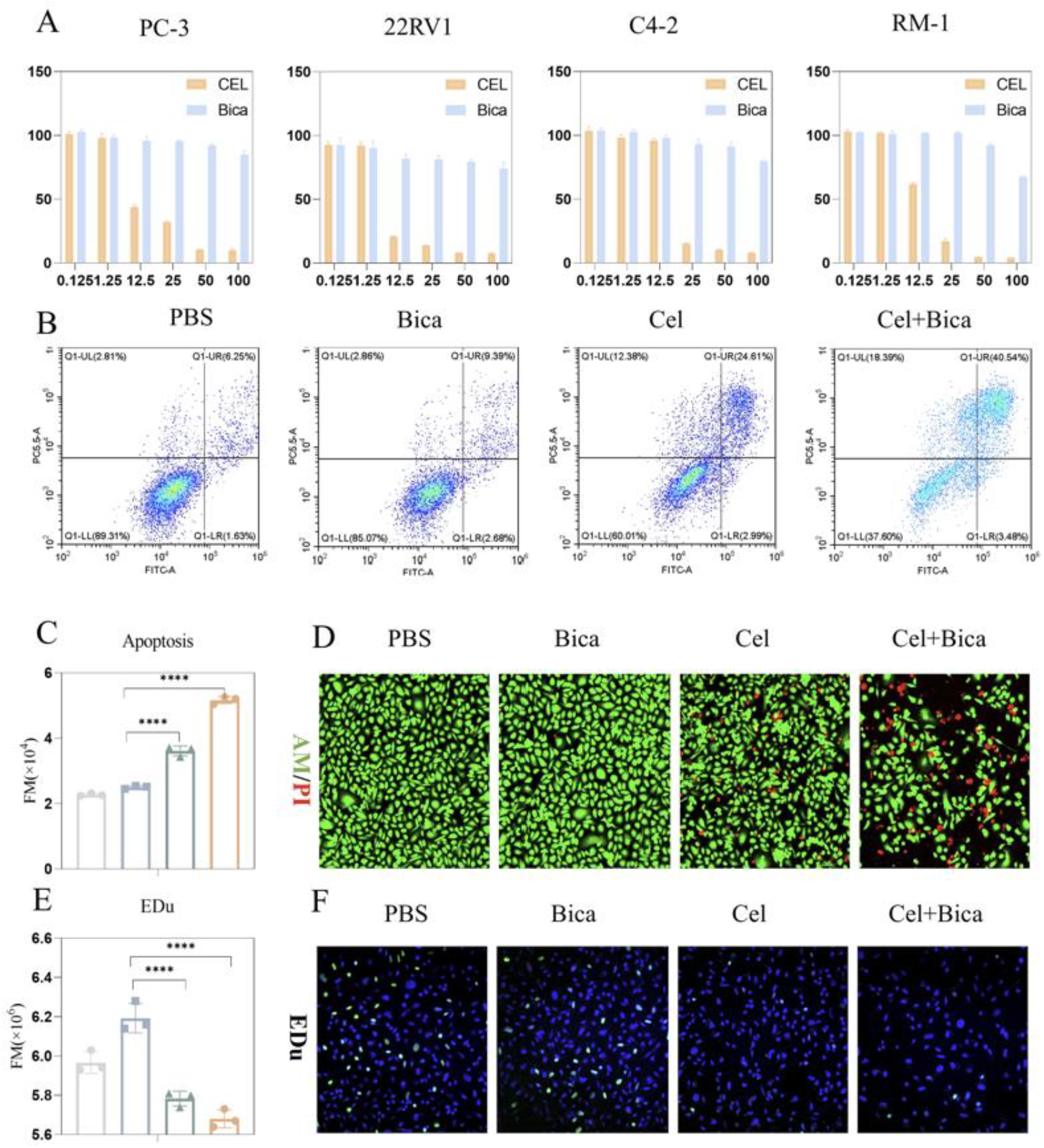
Celastrol enhances the antitumor activity of bicalutamide *in vitro*. (A) Relative Cell viabilities of PC-3, 22RV1, C4-2, and RM-1 Cells with 48 h treatment of Celastrol and bicalutamide via MTT assay, respectively. (B) Representative FCM images and (C) quantification of apoptotic rate via FCM in PC-3 Cells treated with various. (D) The CLSM images of PC-3 Cells stained with calcein-AM (green, viable) and PI (red, dead) after various treatments. (E) FCM images of fluorescence value and (F) the CLSM images of PC-3 Cells stained with Edu. Data are presented as mean ± SD. P values were calculated via one-way ANOVA with Dunnett multiple comparison test. ** *p* < 0.01, **** *p* < 0.0001. **Fig.1 — Source data 1** **PDF file containing original IF Source data for Figure 1D and F, indicating the relevant bands and treatments**.

### 3.2 *In vitro* Celastrol-induced ferroptosis study

Ferroptosis is distinct from traditional forms of cell death, like apoptosis, necrosis, an d autophagy. It mainly occurs due to the accumulation of lipid peroxides that depend on iron, causing mitochondrial contraction and rupturing the mitochondrial membrane, which ultimately leads to cell death (25). We first investigated how Celastrol induces ferroptosis in vitro by examining the mitochondrial membrane potential of tumor cells treated with various drugs using the JC-1 assay kit. On one hand, CLSM results indicated that PC-3 cells treated with PBS exhibited weak green fluorescence, indicating mitochondrial damage, along with prominent red fluorescence, indicating normal mitochondria. In contrast to the bicalutamide (Bica) group, the cells treated with Cel + Bica exhibited almost no red fluorescence and increased green fluorescence, indicating that Cel + Bica treatment resulted in significant mitochondrial damage in PC-3 cells (Figure 2A).

On the other hand, mitochondria are the primary source of reactive oxygen species (ROS) and play a dual role in ferroptosis, serving as a critical site for this process. There is a positive correlation between ROS fluorescence intensity and ferroptosis. Ferroptosis is contributed to by excessive intracellular ROS accumulation, reduced GSH levels, diminished GPX4 scavenging, and an imbalance in ROS scavenging and degradation homeost asis. Furthermore, studies have shown that the generation of a large amount of ROS, can lead to the ICD effect in tumor Cells. Western blot analysis revealed that drug treatment reduced the expression levels of ferroptosis target proteins (GPX4, SLC7A11, and SCD). The protein levels in the Cel and Cel + Bica groups were significantly lower than in the Bica group (Figure 2B). Additionally, statistical analysis of the gray value showed a significant difference in protein content between the Cel and Cel + Bica groups (Figure S1B, 1C). The intraCellular ROS levels in PC-3 Cells were measured using the fluorescent probe H2DCFDA (Figure 2C). The results indicated that the Cel + Bica group exhibited the strongest green fluorescence, suggesting that Cel + Bica had the highest ability to induce ROS (Figure 2C). Subsequently, the ROS levels in Cells treated with each drug were semi-quantified by FCM. The results showed that the Cel + Bica group induced th e highest ROS generation in PC-3 Cells, which was 1.73 times higher than that induced by Bica and 1.36 times higher than that induced by Cel (Figures 2C, 2D).

Furthermore, excessive production or stimulation of reactive oxygen species (ROS) ca n lead to lipid peroxidation, ultimately resulting in tumor cell death. We utilized MDA assay kit to detect the content of MDA in PC-3 Cells after drug treatment. The MDA content after treatment with Cel + Bica was found to be 2.15 times higher than that after Bica treatment. The MDA content after treatment with Cel was found to be 1.84 times higher than that in the non-drug-treated group (Figures 2E). We used the BODIPY™ 581/ 591 C11 fluorescent probe to incubate PC-3 cells and detect lipid peroxidation expression in tumor cells. CLSM observed that the fluorescent expression of Cells treated with Cel and Cel + Bica was stronger than that of Bica treatment (Figures 2F). Flow cytometry revealed that the Cel + Bica group showed the strongest fluorescence expression of lipid peroxidation after drug treatment, being 1.11 times higher than that of the Bica group (Figures 2G).

Ferroptosis is a form of cell death that differs from apoptosis, which is programmed cell death, and necrosis, which is uncontrolled cell death. It is driven by the accumulation of lipid peroxides that depend on iron, leading to mitochondrial damage and cell death. The combination of Celastrol and bicalutamide significantly increased mitochondrial damage and reactive oxygen species (ROS) generation in PC-3 cells. MDA levels rose 2.15 -fold compared to treatment with bcalutamide alone, indicating increased lipid peroxidation and the induction of ferroptosis.

**Fig. 2.**
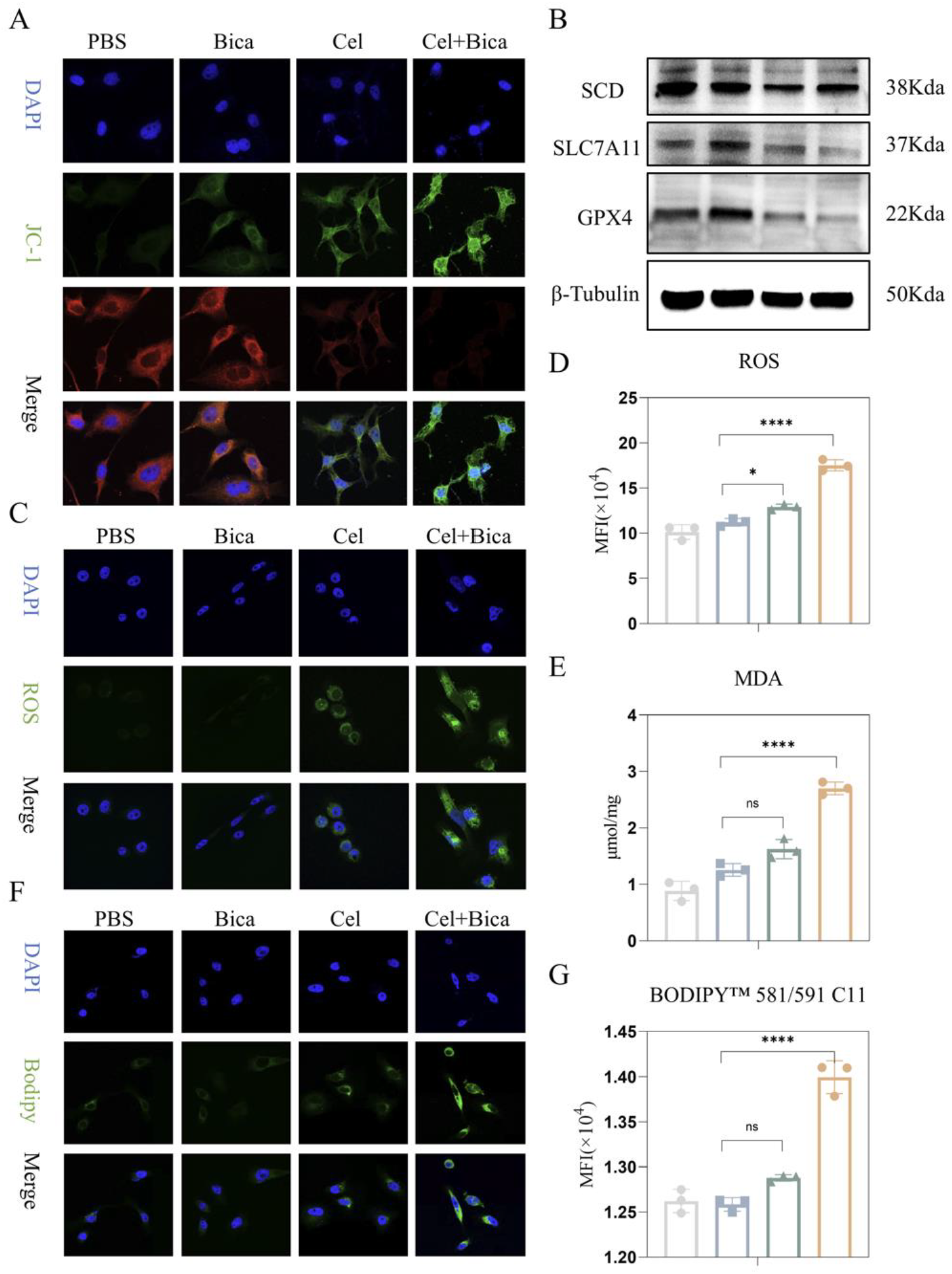
Celastrol-induced ferroptosis of bicalutamide *in vitro*. (A) Representative CLSM images of JC-1 in PC-3 Cells upon various treatments. (B) Western Blot of GPX4, SCD, SLC7A11 and β-Tubulin protein. (C) Representative CLSM images of ROS in PC-3 Cells after various treatments. (C) ROS generation in PC-3 Cells after various treatments by flow cytometry. (E) MDA in PC-3 Cells after various treatments. (F) Representative CLSM images and FCM analysis of lipid peroxidation in PC-3 Cells after various treatments. Data are presented as mean ± SD. P values were calculated via one-way ANOVA with Dunnett multiple comparison test. ** *p* < 0.01, **** *p* < 0.0001. **Fig.2 — Source data 1** **PDF file containing original IF Source data for Figure 2A, C and F, indicating the relevant bands and treatments**. **Fig.2 — Source data 4 and 5** **PDF file containing original western blots data for Figure 2B, indicating the relevant bands and treatments**.

### 3.3 Celastrol promotes anti-tumor immunity *in vitro*

Research indicates that chemotherapy and photodynamic therapy can trigger immunogenic cell death (ICD). The immunogenicity of ICD is primarily mediated by damage-asso ciated molecular patterns (DAMPs). During this process, tumor cells secrete high-mobility group box 1 (HMGB1) from the nucleus, release adenosine triphosphate (ATP) from the cytoplasm, and translocate calreticulin (CRT) to the surface of the tumor cell, which collectively enhances ICD. Antigen-presenting cells recognize these events, which promote I CD.

To investigate the migration pattern of HMGB1 in PC-3 cells, we performed CLSM analysis (Figures 3A, 3E). In comparison to cells treated with Celastrol or bicalutamide alone, the Cel + Bica group showed increased red fluorescence in the cytoplasm and decreased red fluorescence in the nucleus. This suggests that the combination of Celastrol an d bicalutamide promotes the extracellular migration of HMGB1 (Figures 3A, 3E).

Next, we analyzed the ATP levels in PC-3 cells. Figure 3C shows that the Cel + Bi ca group had the lowest ATP concentration at 1.09nM. In comparison, ATP levels in cell s treated with bicalutamide were 6.78nM, and those treated with Celastrol were 1.44nM. Both levels were significantly higher, being 6.22 times and 1.32 times greater, respectively, than the levels in the Cel + Bica group. This indicates that treatment with bicalutamide and Celastrol led to a substantial loss of ATP from the PC-3 cells.

Furthermore, we observed how various drug treatments can induce CRT exposure on the surface of PC-3 cells using CLSM. The results showed that bicalutamide, Celastrol, a nd Cel + Bica treatments all induced CRT expression on the cell surface, with the Cel + Bica group displaying the strongest red fluorescence, as indicated in Figures 3B and 3 D. These results suggest that the ICD effect induced by Cel + Bica treatment is stronger than that of the other treatment groups.

Finally, to confirm the immune activation effect of Celastrol-treated tumor cells, we isolated purified bone marrow-derived dendritic cells (BM-DCs) from mice. We co-culture d various drug-treated PC-3 cells with BM-DCs in a trans-well system for 24 hours. The maturation of BM-DCs was then analyzed semi-quantitatively using flow cytometry (FC M). The results indicated that the BM-DC maturation rate in the Cel + Bica group was 64.3%, higher than the bicalutamide group at 55.1% and the Celastrol group at 74.73% (Figures 3F and 3G). In summary, Celastrol sensitizes bicalutamide, enhances DAMP release from tumor cells, and induces BM-DC maturation, thereby activating ICD.

The results indicate that chemotherapy drugs can induce immunogenic cell death (IC D) via damage-associated molecular patterns (DAMPs). In PC-3 cells, the combination of Celastrol and bicalutamide enhances HMGB1 migration, ATP release, and calreticulin (C RT) surface exposure, which leads to increased dendritic cell maturation compared to treatments with either drug alone.

**Fig. 3.**
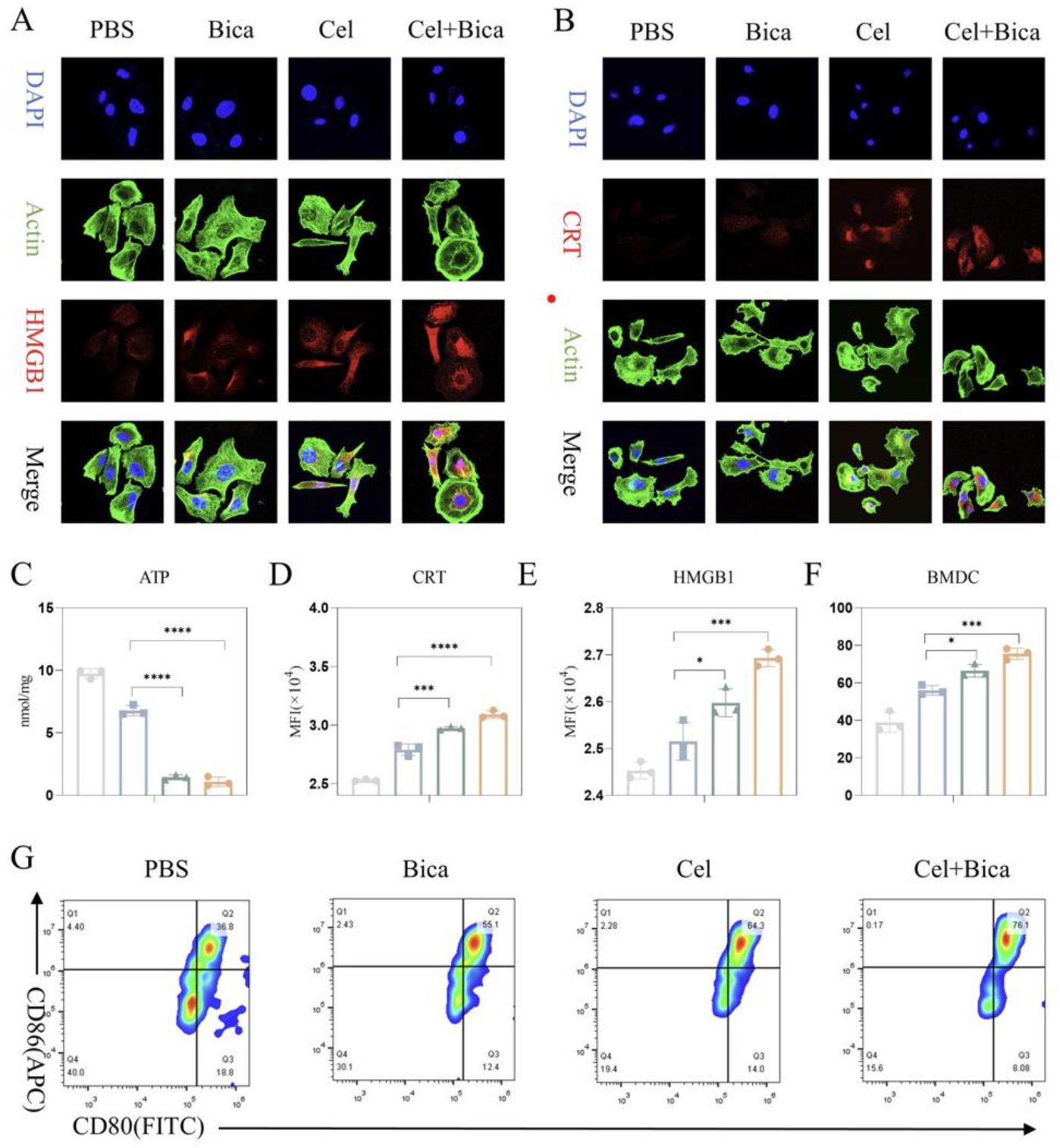
Celastrol promotes anti-tumor immunity *in vitro*. (A) Representative CLSM images analysis and (E) the corresponding quantification of the HMGB1 release in PC-3 Cells after various treatments. (B) Representative CLSM images analysis and (D) the corresponding quantification of CRT in PC-3 Cells upon various treatments. (C) The level of ATP in PC-3 Cells. (F) Representative FCM analysis and (G) quantitative study of DC maturation co-cultured with RM-1 Cells with various treatments. Da ta are presented as mean ± SD. P values were calculated via one-way ANOVA with Dunnett multiple comparison test. ** *p* < 0.01, **** *p* < 0.0001. **Fig.3 — Source data 1** **PDF file containing original IF Source data for Figure 3A and B, indicating the rele vantbands and treatments**.

### 3.4 *In vivo* analysis of antitumor activity

We established a C57BL/6 mouse model of CRPC using RM-1 cells to investigate the anti-tumor effects of Celastrol and bicalutamide in vivo. RM-1 tumor cells were inoculated at a density of 8*10^5^ cells in the right lower limb of C57BL/6 mice, which were t hen divided into four groups after 7-9 days of tumor formation (n = 5) and treated with PBS, bicalutamide (Bica), Celastrol (Cel), and a combination of Celastrol and bicalutamide (Cel + Bica) on days 0, 2, 4, 6, 8, 10, 12, and 14. Tumor volume and weight were monitored, and on day 14, the mice were euthanized. The tumors were weighed and photographed, as shown in Figure 4A. The results, presented in Figures 4C-4D and Figure S 1D, indicated that the PBS, Bica, and Cel groups experienced rapid tumor growth, where as the Cel + Bica group exhibited a significant reduction in the tumor growth rate.

Additionally, we analyzed tumor tissue sections for necrosis and cell apoptosis using Hematoxylin and Eosin (H&E) staining and TUNEL (Terminal deoxynucleotidyl transferase dUTP nick-end labeling). The results indicated that the group treated with Celastrol an d bicalutamide had the highest levels of tumor necrosis and cell apoptosis. This suggests that Celastrol effectively enhances the cytotoxicity of bicalutamide, promotes apoptosis in tumor cells, inhibits tumor proliferation, and increases anti-tumor efficacy (Figure 4E).

**Fig. 4.**
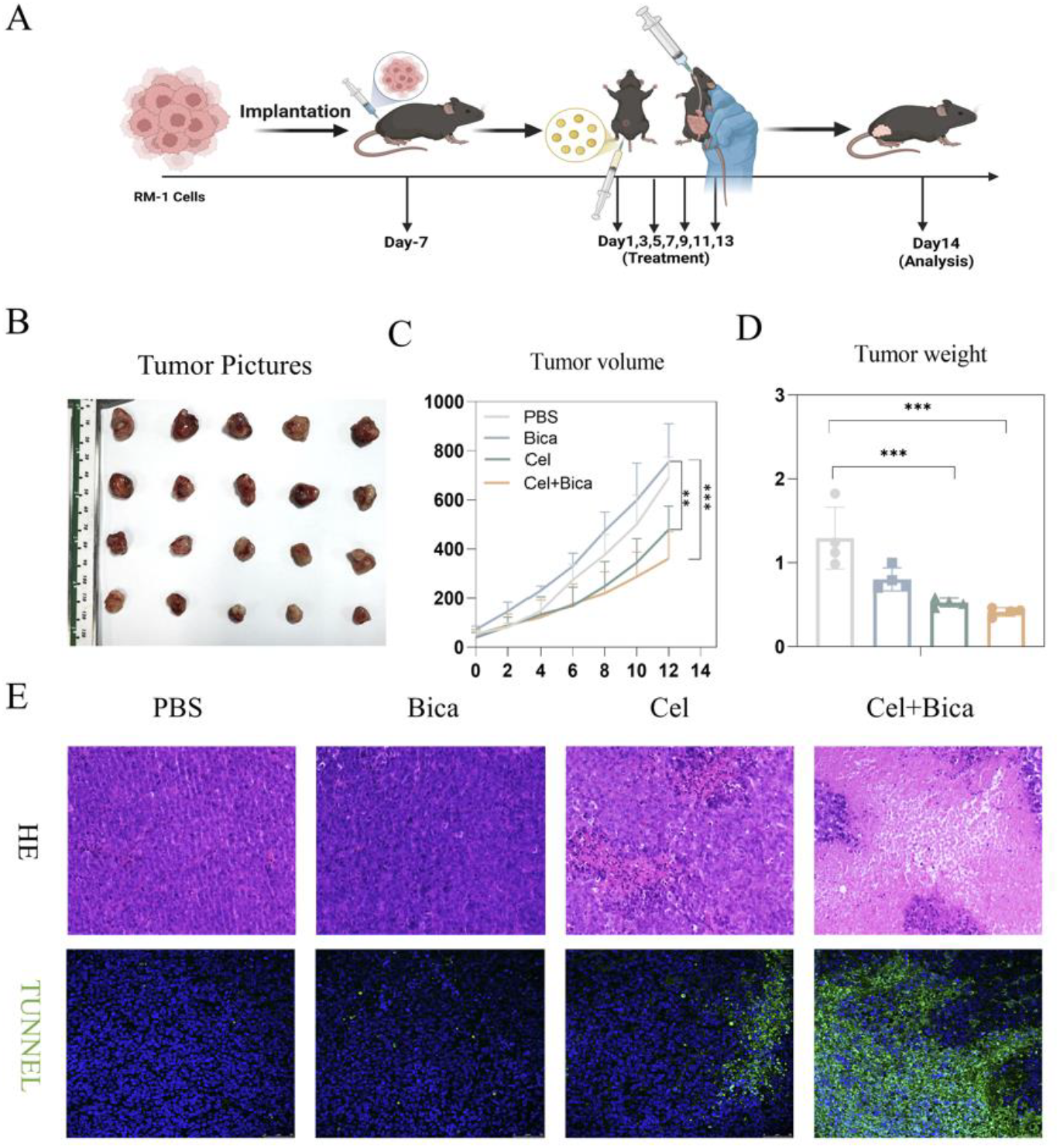
Celastrol enhances the antitumor activity of bicalutamide *in vivo*. (A) Schematic illustration of *in vivo* treatment schedule. (www.biorender.com). (B) Images, (C) volumes, and (D) weights of tumors obtained from mice at day 14 after different treatments. (E) Representative images of H&E and TUNEL staining of tumor tissues. Data are presented as mean ± SD. P values were calculated via one-way ANOVA with Dunnett multiple comparison test. ** *p* < 0.01, **** *p* < 0.0001. **Fig.4 — Source data 3** **PDF file containing original IF Source data for Figure 4E, indicating the relevantban ds and treatments**.

### 3.5 *In vivo* anti-tumor immune study

To investigate the effects of the Celastrol and bicalutamide combination therapy on immune system activation in vivo, we performed immunofluorescence staining. This staining allowed us to assess the infiltration of CD8^+^ T cells, CRT translocation, and HMGB1 migration in tumor tissues (Figure 5A-Figure 5C). The tumor tissues in the Bica and Cel groups showed greater infiltration of CD8^+^ T cells, CRT translocation, and HMGB1 cytoplasmic release compared to the control group. Notably, the Cel + Bica group exhibited significantly stronger fluorescence than the bicalutamide and Celastrol groups (Figure 5A-Figure 5C).

During ICD, tumor cells release DAMPs, which promote the maturation of dendritic cells (DCs) and activate the adaptive immune response. Therefore, we conducted FCM analysis to evaluate the maturation of DCs in tumors from mice treated with various therapies (Figure 5D). The results indicated that the proportion of mature DCs (CD80^+^ CD86^+^) in the Celastrol + bicalutamide (Cel + Bica) group (46.1%) was 1.81 and 1.52 times higher than in the bicalutamide (Bica) group (25.4%) and the Celastrol (Cel) group (30.4%), respectively (Figure 5D). These findings suggest that Celastrol enhances bicalutamide’s effect on promoting DC maturation.

**Fig. 5.**
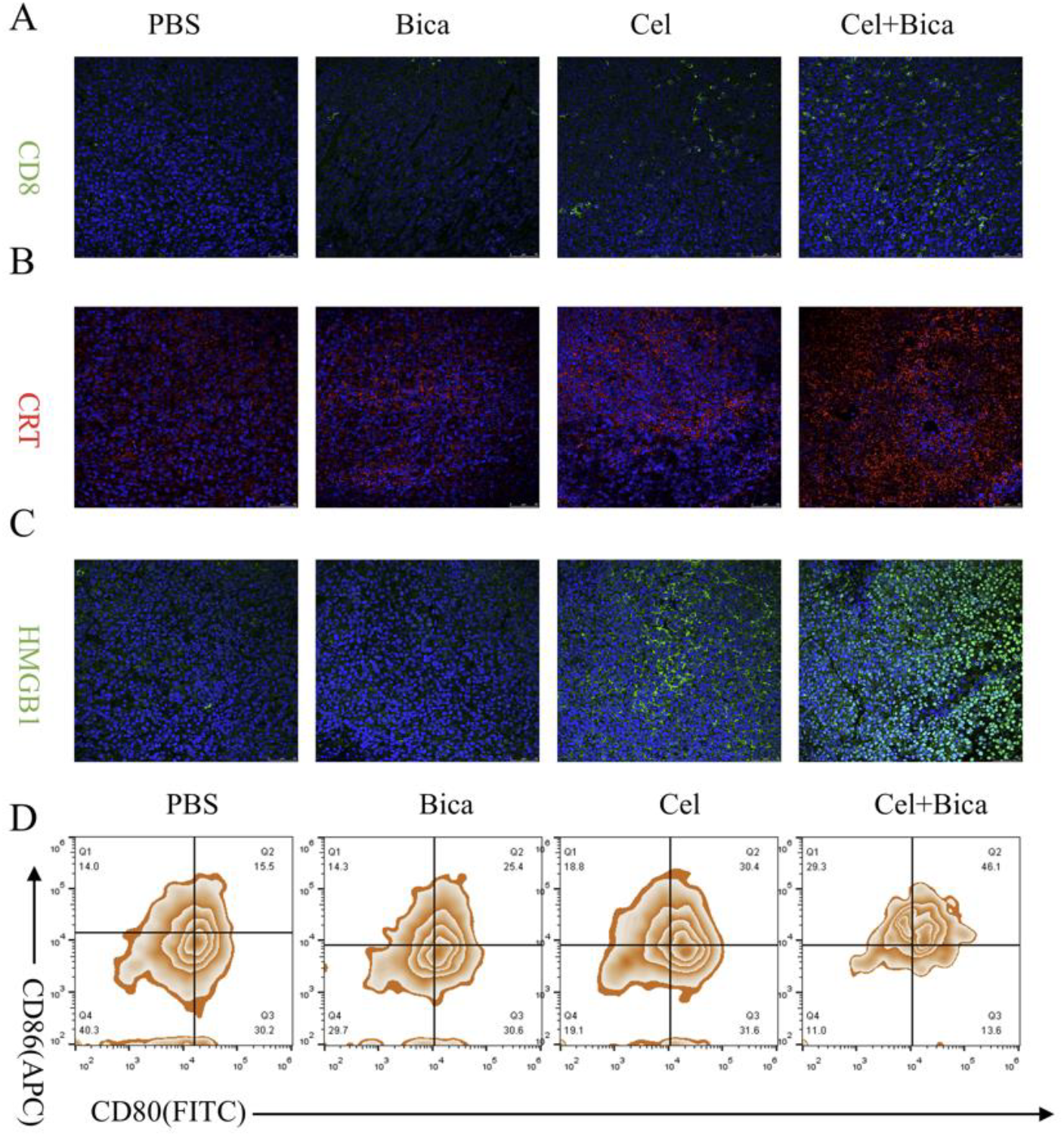
Celastrol enhances the antitumor immune effect of bicalutamide *in vivo*. (A) Representative immunofluorescence images of the infiltration of the CD8^+^ T Cells (above), (B) the expression of CRT (middle), and (C) the release of HMGB1 (below) staining sections within tumor after various treatments. (D) Representative flow cytometric analysis of mature DCs (CD80^+^ CD86^+^) populations within lymph tissues after various treatments. **Fig.5 — Source data 2** **PDF file containing original IF Source data for Figure 5A and B, indicating the rele vantbands and treatments**. **Fig.5 — Source data 3** **PDF file containing original IF Source data for Figure 5C, indicating the relevantban ds and treatments**.

### 3.6 Celastrol activates the tumor immune microenvironment

We collected tumors, spleens, and lymph nodes from C57BL/6 mice treated with different drug regimens to analyze how the Celastrol and bicalutamide combination therapy impacts the immune microenvironment. Relevant immune markers were then characterized by flow cytometry.

First, we analyzed the maturation of DC cells in the lymph and tumor tissues of mice using flow cytometry (Figure 6B, 6G). In the lymph of C57BL/6 mice, the proportion of DC cells in the Cel + Bica group (31.40%) was 1.52 and 1.27 times higher than in the bicalutamide group (20.63%) and Celastrol group (24.67%), respectively (Figure 6B). In the tumor tissue, the proportion of DC cells exceeded that in the lymph, with the C el + Bica group displaying 39.33%, which is higher than the 29.97% in the Bica group and 32.87% in the Cel group. These results suggest that the combination of Celastrol and bicalutamide enhances the maturation of DC cells (Figure 6G).

Mature dendritic cells (DCs) present antigens to T cells in the lymph nodes, activating CD8^+^ T cell responses and starting an effective adaptive immune response. Therefore, t he next analysis focused on the relative proportion of CD8^+^ T Cells in the spleen and tumor tissues (Figure 6A, 6E, 6F). The results showed that in the spleen, the proportion o f CD8+ T cells in the Cel + Bica group (40.03%) was 1.68 and 1.43 times higher than in the Bica group (23.77%) and Cel group (28.07%), respectively (Figure 6 A, 6E). Similar results were observed in the tumor, where the proportion of CD8^+^ T cells in the C el + Bica group (56.13%) was 1.54 and 1.30 times higher than that in the Bica and Cel groups, respectively (Figure 6F).

Subsequently, the relative proportion of MDSC Cells (CD45^+^ CD11b^+^ Gr-1^+^) in the t umor tissue was analyzed. The results revealed that the proportion of MDSC cells in the Cel+Bica group (13.67%) was 2.50 times higher than in the Bica group (5.48%) and 1. 85 times higher than in the Cel group (7.38%) (Figure 6C). Finally, the M1 (CD80^+^ CD 206^−^) and M2 (CD80^−^ CD206^+^) macrophage phenotype ratios were used to assess the eff ectiveness of Celastrol and bicalutamide in initiating anti-tumor immune responses. The r esults indicated that Celastrol and bicalutamide co-treatment induced a shift in tumor-asso ciated TAM Cells from M2 to M1. In particular, the Cel + Bica group exhibited the hig hest M1/M2 ratio in the tumor (1.23), which was 3.85 and 1.70 times higher than that i n the Bica group (0.32) and Cel group (0.72), respectively (Figure 6D). The sentence ca n be made more concise for easier understanding. These findings suggest that co-treatme nt with Celastrol and bicalutamide can reverse the immunosuppressive state of the tumor microenvironment (TME) and enhance the adaptive immune response synergistically.

**Fig. 6.**
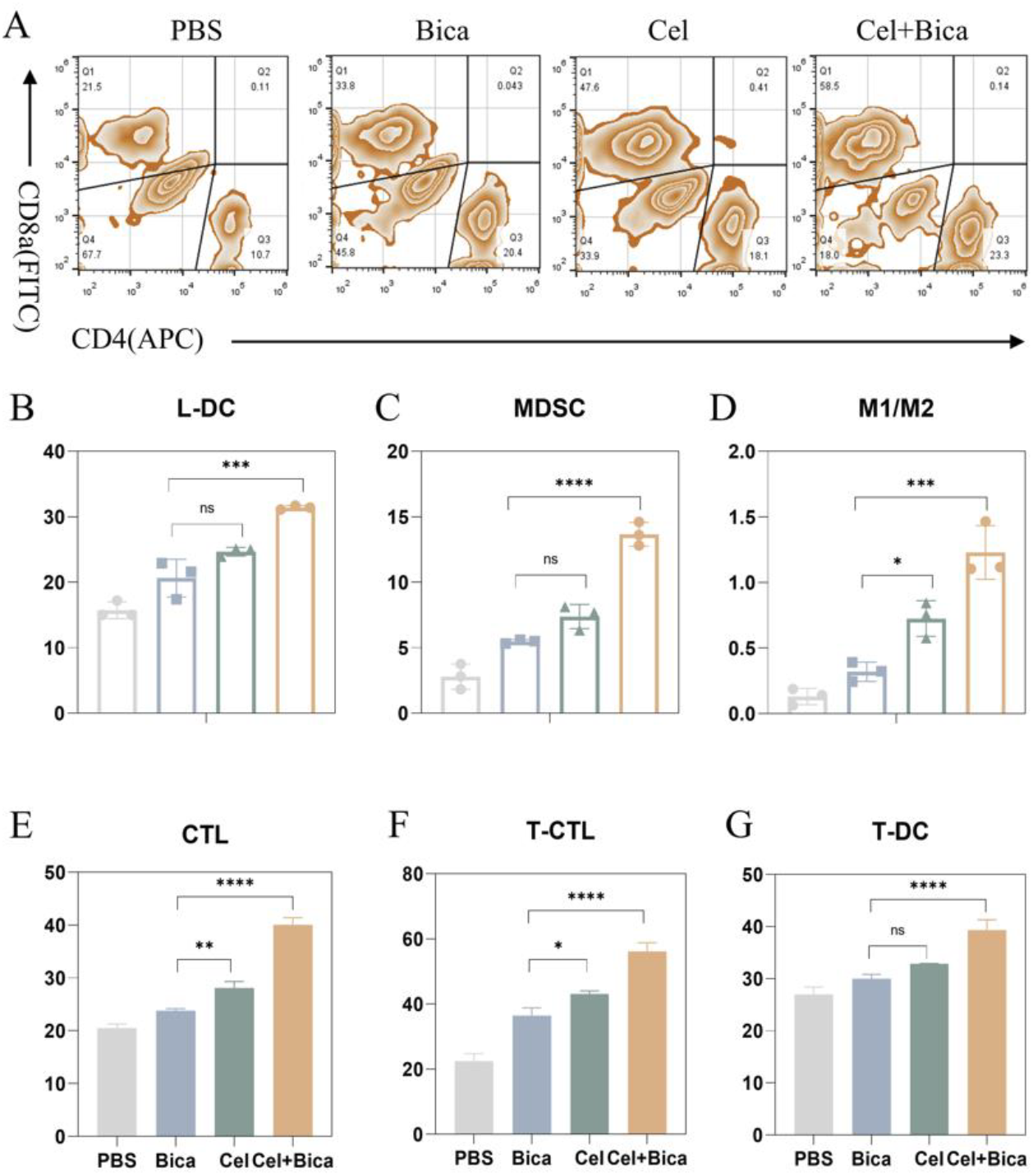
Celastrol enhances the tumor immune microenvironment activation effect of bicalutamide. (A) The percentages of populations of CD8^+^ T Cells within spleen in each group are presented as histograms. (B) The percentages of populations of mature DCs (CD80^+^ CD86^+^) within lymph in each group are presented as histograms. (C) The percentages of populations of MDSCs Cells (CD45^+^ CD11b^+^ Gr-1^+^) within tumor in each group are presented as histograms. (D) The percentages of populations of M1 Cells (CD80^+^ CD206^-^/M2 TAM Cells (CD80^-^ CD206^+^) within tumor in each group are presented as histograms. (E) The percentages of populations of CD8^+^ T Cells within spleen in each group are presented as histograms. (F) The percentages of populations of CD8^+^ T Cells within tumor in each group are presented as histograms. (G) The percentages of populations of mature DCs (CD80^+^ CD86^+^) within tumor in each group are presented as histograms. Data are presented as mean ± SD. P values were calculated via one-way ANOVA with Dunnett multiple comparison test. ** *p* < 0.01, **** *p* < 0.0001.

### 3.7 Studies on the mechanism of ferroptosis *in vivo* and safety of the drug Celastrol

To investigate the in vivo mechanism of Cel and Bica combination therapy, we performed immunofluorescence staining of tumor tissues. By staining for GPX4 and SCD, w e could assess the iron death in the tumor tissues (Figure 7A-figure 7B). The fluorescent staining of GPX4 and SCD in tumor tissues was attenuated in the Cel group compared to the control and Bica groups. Notably, the fluorescence of the Cel + Bica group was significantly weaker than that of the Bica and Cel groups (Figure 7A-figure 7B).

In vivo safety evaluation is a crucial assessment of a drug’s therapeutic efficacy. Cel astrol exhibits some degree of reproductive toxicity. In everyday use, Celastrol is employ ed to treat immune diseases and has recently gained favorable recognition in oncology. H &E staining of the heart, liver, spleen, lungs, and kidneys revealed that cardiac muscle fi bers were neatly aligned, with clear transverse striations and no inflammatory cell infiltration in the interstitium. In all groups, the liver showed structurally intact hepatic lobules, a clear axis of the central vein-portal canal area, and radially arranged hepatocyte cords devoid of fatty vacuoles. In the control and Bica groups, the white and red medulla we re clearly demarcated in the spleen, and there was no dilatation of the marginal zone. The white and red medulla were poorly demarcated in the Cel and Combined groups. In t he lungs, the alveolar structure was intact, there was no thickening of the septa, and there was no hyperplasia of the bronchial epithelium. In kidneys glomerular capillaries were well opened and the brush border of proximal tubules was intact. The above results indicate that the administered dose is safe and the safety of mice is good.

**Fig. 7.**
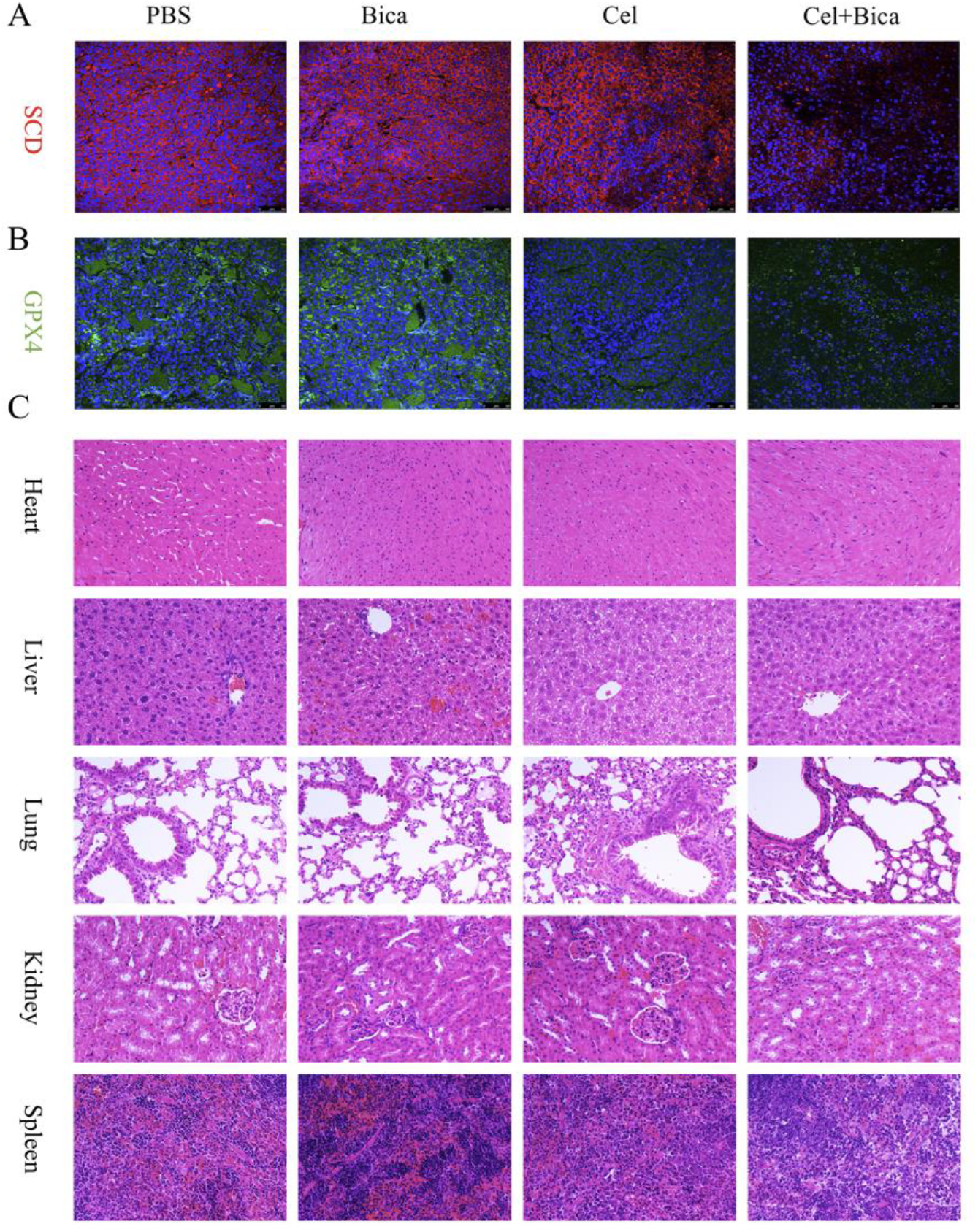
Studies on the mechanism of ferroptosis *in vivo* and safety of the drug Celastrol. (A) Representative immunofluorescence images of GPX4 protein expression staining in tumor sections after different treatments. (B) Representative immunofluorescence images of SCD protein expression staining in tumor sections after different treatments. (C) Representative images of H&E staining of heart, liver, lung, kidney and spleen tissues. Data are presented as mean ± SD. P values were calculated via one-way ANOVA with Dunnett multiple comparison test. ** *p* < 0.01, **** *p* < 0.0001. **Fig.7 — Source data 2** **PDF file containing original IF Source data for Figure 7B, indicating the relevantban ds and treatments**. **Fig.7 — Source data 3** **PDF file containing original IF Source data for Figure 7A, indicating the relevantbands and treatments**.

## 4. Discussion

The present study elucidates a novel mechanism by which Celastrol potentiates bicalutamide’s antitumor efficacy in prostate cancer through dual modulation of ferroptosis and immunogenic cell death (ICD). Our findings advance current understanding of Celastrol’s therapeutic potential and provide critical insights into overcoming bicalutamide resistance.

Celastrol exhibits significant antitumor activity in prostate cancer cell lines, with the highest IC50 in RM-1 cells and enhanced apoptosis when combined with bicalutamide, leading to reduced cell proliferation. Additionally, Celastrol induces ferroptosis in PC-3 cells by causing mitochondrial damage and increasing reactive oxygen species (ROS) levels, as evidenced by elevated MDA content and lipid peroxidation, with the combination treatment showing a 2.15-fold increase in MDA compared to bicalutamide alone.

Celastrol enhances ICD in PC-3 cells by promoting the extracellular migration of H MGB1, reducing ATP levels, and increasing calreticulin (CRT) surface exposure when combined with bicalutamide. This combination treatment leads to a higher maturation rate of bone marrow-derived dendritic cells (BM-DCs), indicating a stronger immune activation compared to treatments with either drug alone.

In vivo analysis of the anti-tumor effects of Celastrol and bicalutamide in a C57BL/ 6 mouse model of CRPC showed that the combination therapy significantly reduced tumor growth and enhanced apoptosis and necrosis compared to individual treatments. Immunofluorescence staining revealed increased CD8^+^ T cell infiltration and dendritic cell maturation in the Cel + Bica group, indicating enhanced immune activation and potential therapeutic efficacy. Safety evaluations indicated that Celastrol is safe at the administered dose, with no significant damage observed in vital organs.

While this work establishes Celastrol/bicalutamide as a promising ferroptosis-ICD co-activator, addressing these limitations and pursuing the outlined directions will clarify its translational potential. Bridging the gap between preclinical efficacy and clinical applicability requires multidimensional validation of both on-target effects and microenvironmental adaptations.

## 5. Conclusion

Celastrol, when used in combination with bicalutamide, induces DC maturation, promotes CD8^+^ T Cell infiltration, CRT ectopia and HMGB1 translocation, and increases the ratio of MDSC and M1/M2 macrophages. These effects significantly activate the immune system and reverse the immune-suppressive microenvironment. As a result, it enhances the cytotoxicity of bicalutamide, promotes tumor Cell apoptosis, inhibits tumor proliferation, and amplifies anti-tumor efficacy. In conclusion, this study demonstrates that Celastrol c an synergistically activate systemic anti-tumor immunity, enhancing the tumor-killing effect of bicalutamide, offering a promising new therapeutic strategy for the clinical personalized immunotherapy of castration-resistant prostate cancer.

## Acknowledgements

The authors are grateful to Dr. He Li and Dong Li for their help with the preparation of figures in this paper. This research was partially supported by Beijing Municipal Natural Science Foundation (grant no. BFHHQS20240003).

## Authors’ contributions

Conceptualization, Peipei Meng and Weibin Yang; methodology, Peipei Meng and Hongdong Huang; software, He Li; validation, Guancheng Wang; formal analysis, Peipei Me ng and He Li; investigation, Yuhang Meng and Dong Li; data curation, Peipei Meng and Qinxin Zhao; writing—original draft preparation, Peipei Meng and He Li; writing—review and editing, Yuhang Meng and Dong Li and Guancheng Wang; visualization, Qinxin Zhao; supervision, Qinxin Zhao and Weibin Yang; project administration, Qinxin Zhao and Hongdong Huang; funding acquisition, Qinxin Zhao. All authors have read and agreed to the published version of the manuscript.

## Data availability statements

The data supporting this article have been included as part of the Supplementary Information.

